# Starch synthesis in *Brachypodium distachyon* endosperm occurs in dynamic, connected amyloplast compartments

**DOI:** 10.1101/2025.09.16.676639

**Authors:** Lara Esch, Qi Yang Ngai, Sergio G. Lopez, Elaine Barclay, Inmaculada Hernández-Pinzón, Matthew J. Moscou, David Seung

**Author notes:** Correspondence: Lara Esch or David Seung.

## Abstract

The morphology of starch granules is a major determinant of the functional and nutritional properties of starch and is highly variable among cereal species. Much of this morphological variation stems from differences in the spatial and temporal patterns of starch granule initiation in amyloplasts during grain development. Simple granules are thought to arise from a single initiation per amyloplast (e.g., in *Brachypodium distachyon*), whereas compound granules develop from multiple initiations per amyloplast (e.g., in rice). Using live-cell imaging in transgenic *Brachypodium* lines expressing a fluorescent amyloplast reporter construct, we discovered that simple-type starch granules in *Brachypodium* can arise from multiple initiations per amyloplast. The amyloplasts showed dynamic changes in their structure, the formation of stromules, and the movement of both the amyloplast within the cell and the starch granules within the amyloplasts. Our results suggest complex and pleiomorphic amyloplast organisation and mobility that can influence starch granule formation and morphology. This goes beyond the existing ‘one granule, one amyloplast’ model for simple-type granules, and advances our understanding of both amyloplast biogenesis and granule formation.

## Introduction

Starch in non-photosynthetic tissues is synthesised in plastid organelles called amyloplasts (Jarvis and López-Juez, 2013). Almost all commercially important starches are produced in amyloplasts of heterotrophic crop organs, such as seeds, tubers and storage organs. Despite this fundamental importance of amyloplasts, very little is known about amyloplast structure and biology.

Amyloplasts play an important role in determining starch granule morphology in the cereal endosperm. Granule morphology among cereal species can be classed into three different categories – simple, compound and bimodal. Simple granules grow from a single initiation per amyloplast, and can be found in species like maize and *Brachypodium* (Myers et al., 2011; Tanackovic et al., 2014; Watson-Lazowski et al., 2022). Compound granules arise from the initiation of multiple granules per amyloplast, which later form a larger, tessellated structure (Matsushima et al., 2013; Matsushima, 2015). The individual granulae contained within compound granules are polygonal, resulting from impaction of individual growing granules within the confines of the amyloplast (Matsushima et al., 2015). Bimodal granules are found exclusively in the Triticeae, where two distinct types of starch granules form. Large A-type granules are initiated and formed in the main amyloplast body during early grain development, whereas small B-type granules initiate later in grain development, partially in protrusions/stromules that emanate from the amyloplasts (Parker, 1985; Langeveld et al., 2000; Matsushima and Hisano, 2019; Esch et al., 2023). The formation of these different granule classes is therefore primarily determined by the spatial and temporal pattern of granule initiation during grain development, where amyloplasts provide the spatial context in which granule initiations can occur.

The influence of amyloplast structure on starch structure has also been demonstrated through genetic mutants. Firstly, mutations that alter amyloplast membrane properties affect starch morphology. The *opaque5* (*o5*) mutant of maize is defective in the monogalactosyldiacylglycerol synthase necessary for normal galactolipid production in amyloplast membranes, and has altered starch granule structure, including the formation of some compound starch granules separated by amyloplast membrane (Myers et al., 2011). Also, the maize *Ven1* mutant defective in a β-carotene hydroxylase has altered carotenoid and membrane composition in amyloplasts, giving rise to a persistent amyloplast envelope and altered starch granule morphology (Wang et al., 2020). Secondly, mutations that alter amyloplast size and shape also affect starch granule morphology. In rice, the *arc5* mutant defective in a plastid division component formed irregular, pleiomorphic amyloplasts, and the individual granulae within compound granules were smaller than those of wild-type, and many had irregular morphology (Yun and Kawagoe, 2009). In wheat, the mutation of the plastid division component, PARC6, resulted in large amyloplasts and striking deformations in A-type granule morphology and altered starch granule size distribution (Esch et al., 2023). The influence of plastid membranes on starch granule structure is not only limited to envelope membranes, but also internal membranes. Arabidopsis mutants with disrupted thylakoid architecture have altered morphology of stromal pockets in between thylakoids where starch granules form, and consequent alterations in starch granule size and shape (Esch et al., 2022; Ilse et al., 2025).

In cereal endosperms, amyloplasts differentiate from proplastids early during grain development, and have a double membrane envelope that is typical of plastids (Buttrose, 1960). Unlike chloroplasts, they have limited internal membrane structures, although there have been reports of tubule-like invaginations of the inner envelope membrane (Buttrose, 1960; Buttrose, 1963). Studying amyloplast structure in the endosperm is technically challenging because at later stages of grain development, the envelope contacts the growing granule and cannot be distinguished from the border of the starch granule under electron microscopy. The envelope only becomes visible when there is free stromal space around the edges of the granule, or when there is stromule formation (Hawkins et al., 2021; Chen et al., 2024). Recently, the use of genetically encoded markers targeting a fluorescent protein to the amyloplast stroma enabled the study of amyloplast morphology using live cell imaging. This approach has been carried out in both barley and rice, where amyloplasts in developing grains were observed to have an elongated structure and contain multiple starch granules (or in the case of rice, multiple compound starch granules), with constrictions between the starch granules (Matsushima and Hisano, 2019). More recently, a similar approach was used to study amyloplast division in Arabidopsis ovule integuments, a non-photosynthetic tissue where most amyloplasts contain multiple granules, resembling compound granules of cereals (Fujiwara et al., 2024). The presence of stromules/protrusions that appear to connect amyloplasts have been observed in both barley endosperm and Arabidopsis ovule integuments (Matsushima and Hisano, 2019; Fujiwara et al., 2024). In the latter, the stromules were proposed to play an important role in one of the modes of amyloplast replication (“stromule-mediated fission”)(Fujiwara et al., 2024).

This live-cell approach has been applied to study amyloplasts in species containing bimodal and compound starch granules, but not yet in a species containing simple starch granules. However, many economically important starches are simple granules (e.g., maize, potato, some yams)(Chen et al., 2021). We therefore aimed to study amyloplast structure and dynamics in *Brachypodium distachyon*, using an amyloplast marker to investigate how changes in amyloplast structure facilitate the synthesis of simple starch granules. *Brachypodium* endosperm is advantageous for the cell biological analysis of amyloplasts as its starch content is not as high as cultivated cereals like wheat and barley, such that its endosperm cells do not become overcrowded with starch granules (Trafford et al., 2013; Watson-Lazowski et al., 2022). Rather than the observation of many discrete amyloplasts, we provide evidence that amyloplasts form a network of connected compartments, many of which contained multiple simple-type starch granules. Connections were through long-lived protrusions that allow stromal exchange between compartments, which were distinct from short-lived, stromule-like protrusions. Further, we reveal amyloplast movement in cytoplasmic streaming as a previously unconsidered physical force that could influence granule morphology. Overall, our study reveals more complexity to amyloplast structure in species producing simple-type starch granules than once thought.

## Results

### Plants expressing a transgenic amyloplast reporter show a normal growth phenotype

To investigate the influence of amyloplast morphology and dynamics on the size and shape of simple-type starch granules in developing *Brachypodium distachyon* endosperm, we created transgenic *Brachypodium* lines expressing an amyloplast reporter construct, which enabled visualisation of amyloplasts using live-cell fluorescence microscopy.

The construct encoded an mCherry fluorescent protein fused to the rice *Granule Bound Starch Synthase* (*GBSS*) chloroplast transit peptide (cTP) at its N-terminus (Fig. 1a), which allowed targeting of mCherry to the amyloplast stroma (Esch et al., 2023). Transgene expression was driven under the constitutive *maize ubiquitin* promoter. We selected a line with robust mCherry expression for further experiments. The transgenic plants stably expressing the reporter protein (Fig. S1) showed a similar growth phenotype and grain size to the wild type (Fig. S1), suggesting that the fluorescent protein did not obviously impact plant development and physiology.

**Figure 1:**
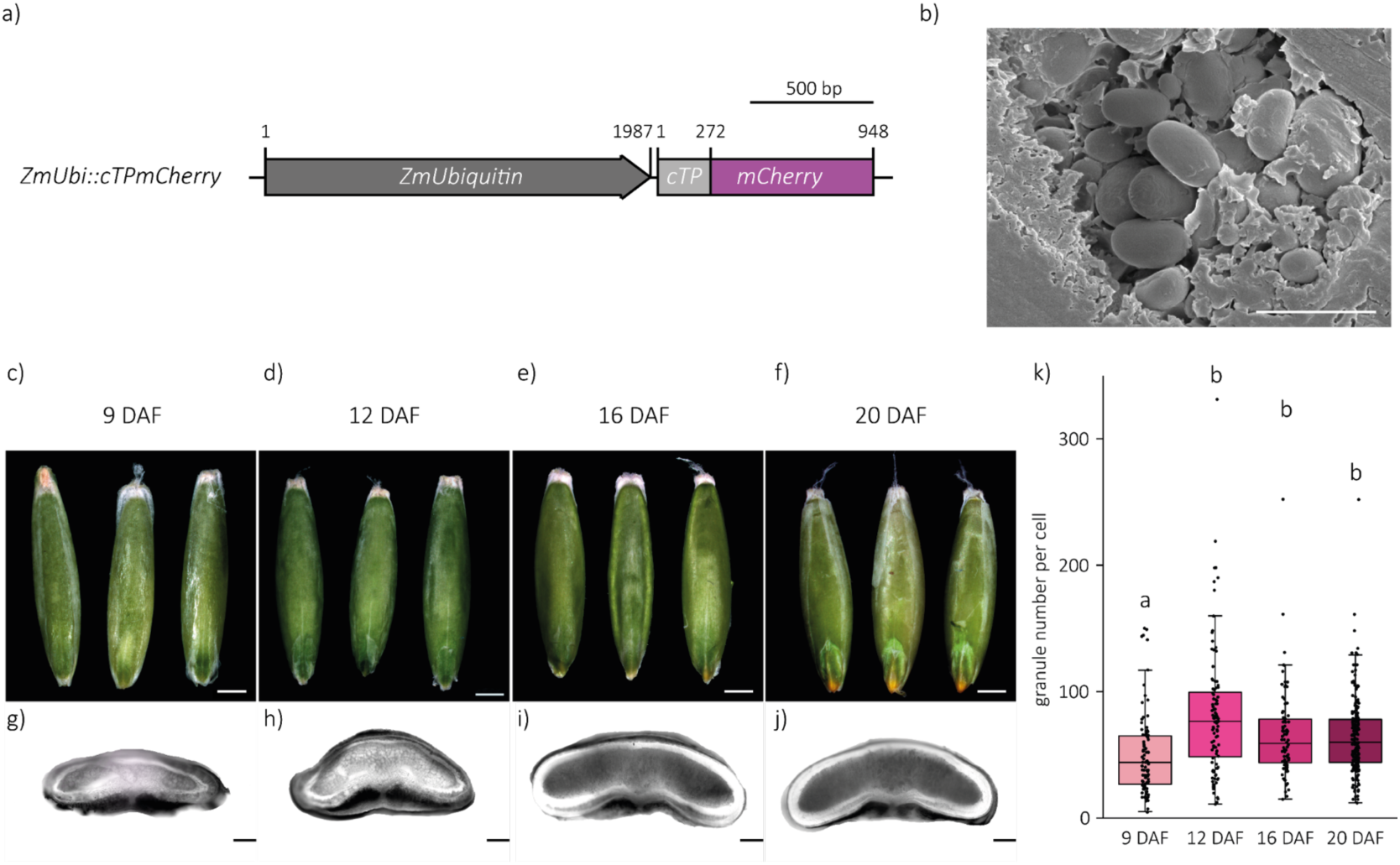
Growth phenotype of *Brachypodium distachyon* amyloplast reporter lines. (a) Schematic illustration of the amyloplast reporter construct consisting of a N-terminal rice Granule Bound Starch Synthase (GBSS) chloroplast transit peptide (cTP) and a codon optimised mCherry driven by a maize Ubiquitin promoter (*Zm*Ubi). (b) Scannning electron micrograph (SEM) of *Brachypodium* starch granules in a cross section through a mature grain. Bar = 5 µm. (c-f) Photographs of 3 representative *Brachypodium* Bd21 seeds overexpressing the amyloplast reporter construct at 9, 12, 16 and 20 days after flowering (DAF). (g-j) Stitched light microscopy images of cross sections of representative *Brachypodium* Bd21 seeds at 9, 12, 16 and 20 DAF. (k) Starch granule number per cell during endosperm development in *Brachypodium* Bd21 ZmUbi:cTPmCherry seeds at 9, 12, 16 and 20 DAF. Starch granule number was quantified in 84-203 cells per time point. Significant differences between the genotypes under a Kruskal– Wallis one-way ANOVA on ranks and all pairwise multiple comparison procedures (Dunn’s method) are indicated with letters, where different letters represent significant differences (P < 0.001).

In the endosperm, *Brachypodium* forms simple-type starch granules. These granules have an ovoid shape and vary in size in the mature grain (Fig. 1b). Starch content in *Brachypodium* is known to increase steadily during grain development, but it is unknown how this relates to changes in granule number (Guillon et al., 2012; Trafford et al., 2013). We therefore quantified the number of granules per endosperm cell during development (9, 12, 16, 20 Days After Flowering - DAF) in the amyloplast reporter lines (Fig. 1c-j). At 9 DAF, endosperm cells contained an average of 50 granules. This was significantly lower than the numbers counted at the later timepoints, when the average number of granules was between 65-82 (Fig. 1k). Although the granule number at 12 DAF appeared higher than at 16 and 20 DAF, there were no significant differences between these three later timepoints, due to the high variation in starch granule number between the analysed cells (Fig. 1k). Overall, it appears that the number of starch granules stabilised after 12 DAF, suggesting that the initiation of new starch granules in *Brachypodium* mostly occurs before this timepoint.

### Amyloplasts in *Brachypodium distachyon* endosperm contain more than one simple-type starch granule

It was hypothesised that simple-type starch granules form from an individual starch granule initiated per amyloplast during endosperm development, allowing them to grow to their characteristic rounded granule shape (Myers et al., 2011; Seung and Smith, 2019). Using live-cell confocal imaging of the amyloplast reporter line, we visualised amyloplasts and starch granules during grain development. Surprisingly, we observed multiple examples of amyloplasts that contained more than one starch granule at three different developmental stages - 9, 12 and 16 DAF (Fig. 2). The numbers of starch granules per amyloplast in all stages ranged from 1 to 4 (Fig. 2). Some amyloplasts contained multiple granules of similar size (Fig. 2a, b, c, f, h, i) while others contained a mix of larger and smaller granules (Fig. 2d, e, g).

**Figure 2:**
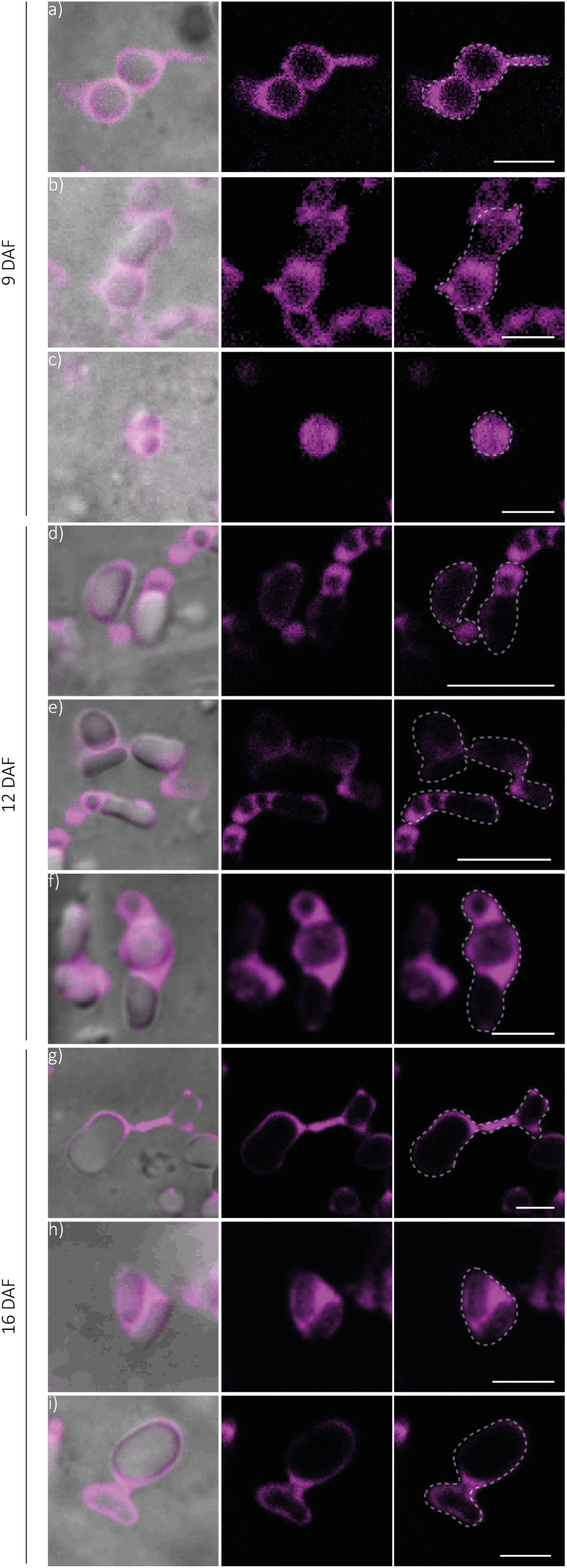
*Brachypodium distachyon* amyloplasts contain multiple starch granules. (a-f) Confocal laser-scanning micrographs of endosperm sections of developing grain at 9, 12 and 16 days after flowering (DAF), in lines stably expressing the amyloplast marker cTPmCherry, shown in magenta. Dashed lines indicate individual amyloplasts containing multiple starch granules. Bars = 3 µm.

The amyloplasts were highly pleomorphic. Those containing only small starch granules appeared round, with stromal space around the granules (Fig. 2c). In most cases, however, they contained large starch granules, and the amyloplast membrane closely enveloped the starch granules, such that organelle’s outline followed the shapes of the granules within. This gave rise to shapes resembling budding yeast (Fig. 2d) or vermiform structures (Fig. 2f). Additionally, we observed protrusions that connected different starch-containing amyloplast compartments (Fig. 2g). To assess the degree to which these structures were influenced by the cellular environment, we observed the morphology of isolated amyloplasts. Amyloplasts were prepared from 16 DAF grains of the reporter line, and the fluorescence of the reporter was used to determine that the amyloplasts were intact. The isolated amyloplasts contained single or multiple starch granules that were variable in size (Fig. 3a-l), and showed the same structural heterogeneity as those in the intact tissues: including budding or vermiform structures (Fig. 3g-j), and even stromules (Fig. 3m-n).

**Figure 3:**
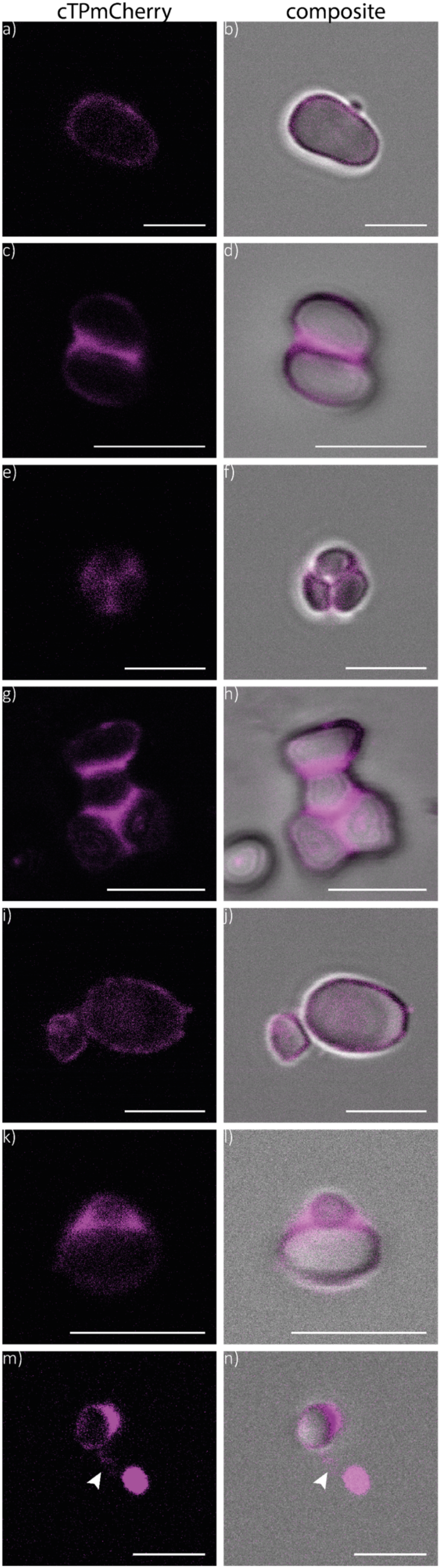
Isolated *Brachypodium distachyon* amyloplasts contain multiple starch granules. (a-n) Confocal laser-scanning micrographs of isolated amyloplasts from developing endosperm at 16 days after flowering (DAF) of lines stably expressing the amyloplast marker cTPmCherry, shown in magenta. Arrow indicates an amyloplast protrusion. We performed three independent amyloplast isolations, all showing similar results. Selected, representative images are presented. Bars = 5 µm.

To confirm these observations in a wild-type background that did not contain the fluorescent reporter, we used Transmission Electron Microscopy (TEM) to observe endosperm amyloplasts in 16 DAF seeds. Similar to our observations in live cells and isolated amyloplasts, many amyloplasts contained multiple granules (Fig. 4a-f).

**Figure 4:**
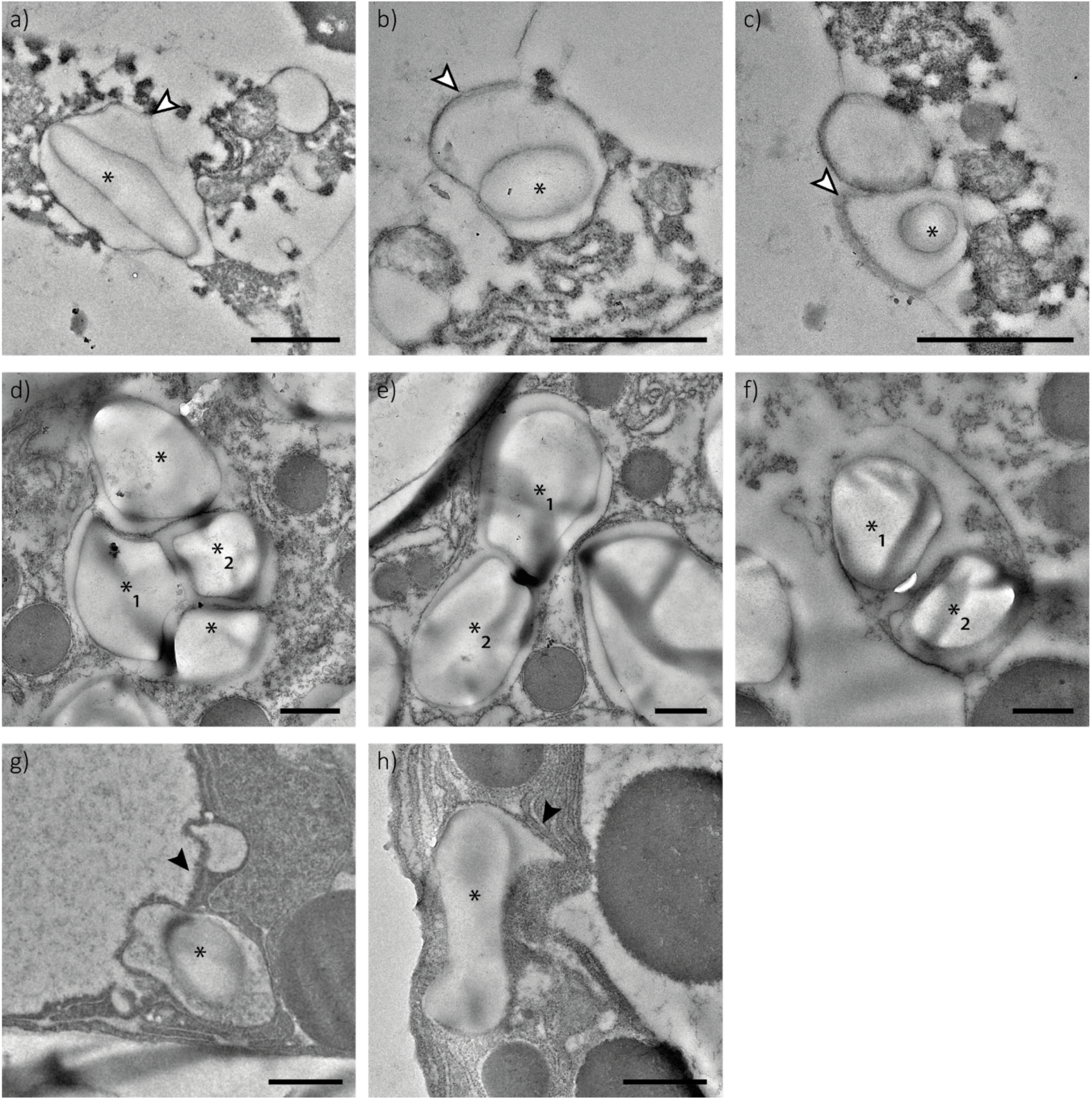
Amyloplast structure in developing grains of *Brachypodium distachyon*. (a-h) Transmission electron microscopy (TEM) images of endosperm sections of a wild-type developing grain at 16 days after flowering (DAF). White arrows indicate the amyloplast membrane, black arrows indicate amyloplast protrusions, and asterisks indicate starch granules. Numbered asterisks indicate multiple starch granules contained within one amyloplast. Bars = 1 µm.

Amyloplasts containing small starch granules were irregular in shape and contained large stromal space between the granule and the membrane (Fig. 4a-c). Amyloplasts with larger granules had envelope membranes tightly associated with the granules, following the contours of the granules, and suggesting that the granule shape influences the amyloplast shape (Fig. 4d-f). Examples of amyloplast protrusions were also visible with TEM, suggesting they were not induced by reporter expression (Fig. 4g, h).

Overall, our results demonstrate that endosperm amyloplasts in *Brachypodium* throughout grain development contain multiple simple-type starch granules, are variable in shape and size, and form protrusions connecting different starch granule containing compartments. These observations strongly contrast the existing hypothesis that simple starch granules arise from one granule initiation event per amyloplast, which would allow the starch granule to grow into a round shape without obstruction from other granules in the plastid. Our observations raise an important question as to how starch granules in *Brachypodium* retain a simple shape despite there being multiple granules in one amyloplast, rather than the multiple granules merging into a compound granule.

### Amyloplasts in *Brachypodium distachyon* endosperm form two types of protrusions

Plastid protrusions have been described in many instances (Hanson and Hines, 2018). Using live-cell imaging, we distinguished two different types of protrusions in *Brachypodium* endosperm amyloplasts of the reporter line. Firstly, we frequently observed the occurrence of stable, long-lived protrusions that connected amyloplast compartments (which we termed ‘type I’ protrusions) (Fig. 5a-f). These protrusions persisted for more than 60 s in the representative example (Fig. 5a-f, Video 1). To test if there is stromal continuum between the connected compartments, we performed Fluorescence Recovery After Photobleaching (FRAP) experiments. Photobleaching one of the interconnected compartments in amyloplasts of 9 and 16 DAF grains resulted in complete bleaching of multiple adjacent compartments, suggesting a stromal continuum between those compartments (Fig. 5g-k and l-p, Fig. S2). However, we did not see fluorescence recovery in the bleached amyloplast compartment, suggesting that the pulse could have bleached all the mCherry protein contained within the connected amyloplast compartments. To test whether that was plausible, we performed Fluorescence Correlation Spectroscopy (FCS) to estimate the diffusion speed of the mCherry protein in the amyloplast stroma. Global fitting of the autocorrelation curves with the triplet state diffusion model determined a diffusion coefficient of 37.650±6.140 µm²/s for the mCherry protein in the amyloplast stroma (Fig. 5q). Using the diffusion coefficient, we determined the mean displacement (MD) of mCherry protein in the amyloplast stroma as 33.31 µm within the span of the 5 s bleaching pulse. Since the distance between the examined amyloplast compartments ranged between 2.35-12.20 µm (Fig. 5 and Fig. S2), it is plausible that all mCherry protein within the connected compartments diffused through the bleaching region within the time of bleaching pulse. Overall, our results clearly demonstrate that there is a stromal continuum between the amyloplast compartments that are connected through type I protrusions.

**Figure 5:**
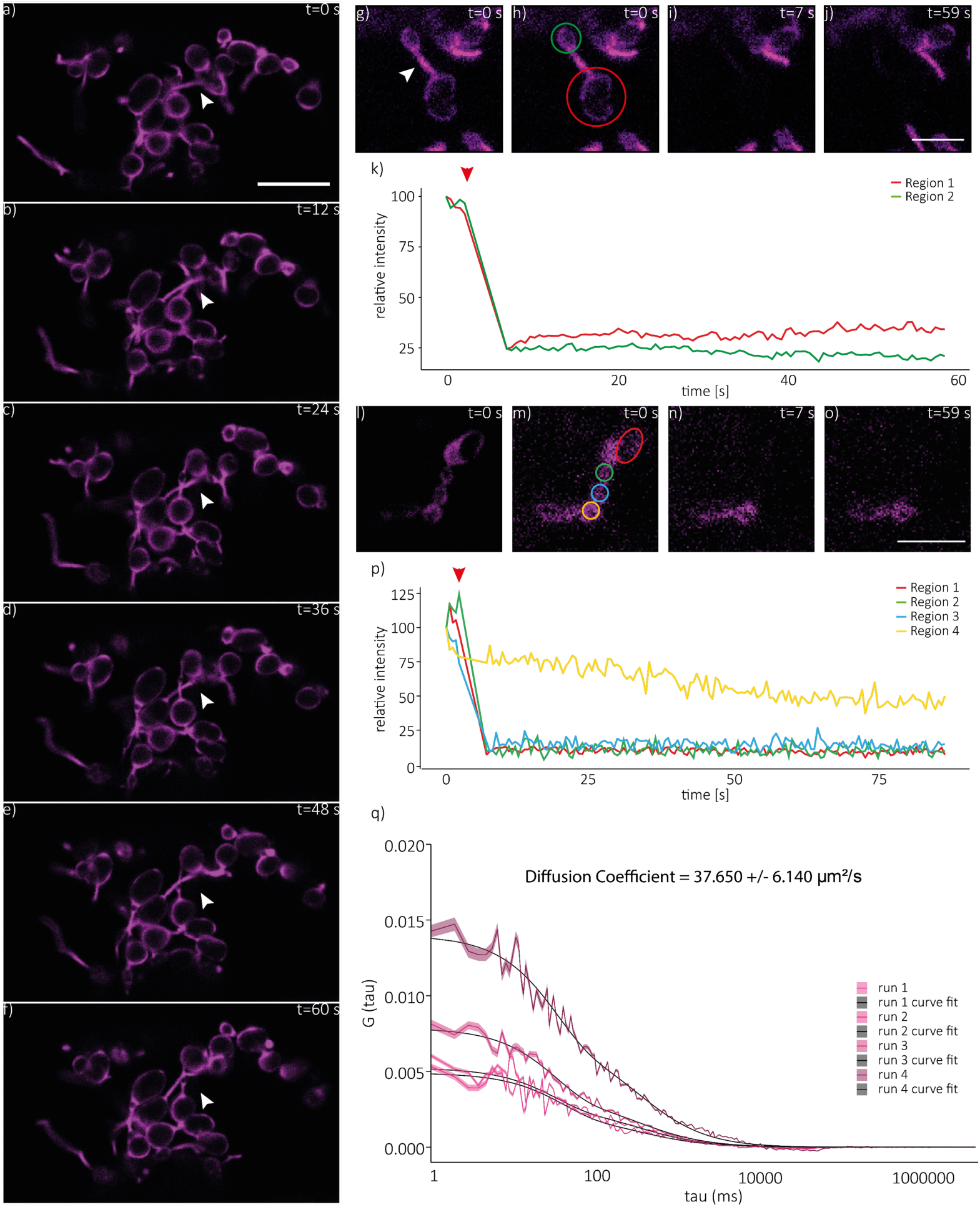
Amyloplasts in *Brachypodium distachyon* endosperm form protrusions that connect amyloplast compartments. Amyloplasts were imaged using confocal laser-scanning microscopy in the reporter lines stably overexpressing the amyloplast marker cTPmCherry, shown in magenta. In all panels, time (t) is shown in seconds (s) (a-f) Time course imaging of developing endosperm sections at 16 days after flowering (DAF). Arrows indicate stable type I protrusions. Bar = 10 µm (g-j) Connected amyloplast compartments in endosperm sections of untreated, developing grain at 16 DAF, before bleaching (g,h) and after bleaching (I,j). Arrow indicates a type I protrusion, red circle indicates region of bleaching and fluorescence intensity measurement (Region 1), and green circle indicates connected region for fluorescence intensity measurement (Region 2). Bar = 5 µm. (k) Relative fluorescence intensity after bleaching measured in regions indicated in (h). Red arrow indicates the timepoint of the bleaching pulse. (l-o) Connected amyloplast compartments in endosperm sections of developing grains treated with 25 µM Latrunculin B at 9 DAF, before bleaching (l,m) and after bleaching (n,o). Red circle indicates region of bleaching and fluorescence intensity measurement (Region 1), coloured circles indicate adjacent regions for fluorescence intensity measurement (Regions 2-4). Bar: 5 µm. (p) Relative fluorescence intensity after bleaching measured in regions indicated in (m). Red arrow indicates timepoint of bleaching pulse. (q) Fluorescence correlation spectroscopy (FCS) curves of mCherry in the stroma. Run data is shown in magenta, and grey lines show curve fitting to the model of diffusion with triplet. Shading represents standard error. Diffusion coefficient was determined using global fit of diffusion with triplet to curves.

In addition to the more stable type I amyloplast protrusions, we observed short-lived protrusions that rapidly extended, moved and retracted, and were not connected to other granule-containing compartments (termed ‘type II’ protrusions) (Fig. 6a). Figure 6 shows a representative time series, showing three different type II protrusions extending, moving, and retracting within a 45 second time frame (Fig. 6a-r, Video 2 and 3). The minimum protrusion length ranged from 0.40 to 2.81 µm, and they reached a maximal length between 4.65-8.52 µm within the 45 s of the time series. To record the movement (including extension and retraction) of the protrusions, we tracked the tip of the protrusion using the Fiji plug-in TrackMate (Ershov et al., 2022) and measured a total travelled distance of 11.1, 6.5 and 11.4 µm for the 3 protrusions respectively within the 45 s time series (Fig. 6s). The maximal distance travelled per frame ranged from 2.2 to 5.6 µm, resulting in a maximum travel speed of 0.26 to 0.82 µm per second. Over the observed 45 s time period, the mean travel speed for the protrusions was 0.13, 0.08 and 0.23 µm/sec, respectively.

**Figure 6:**
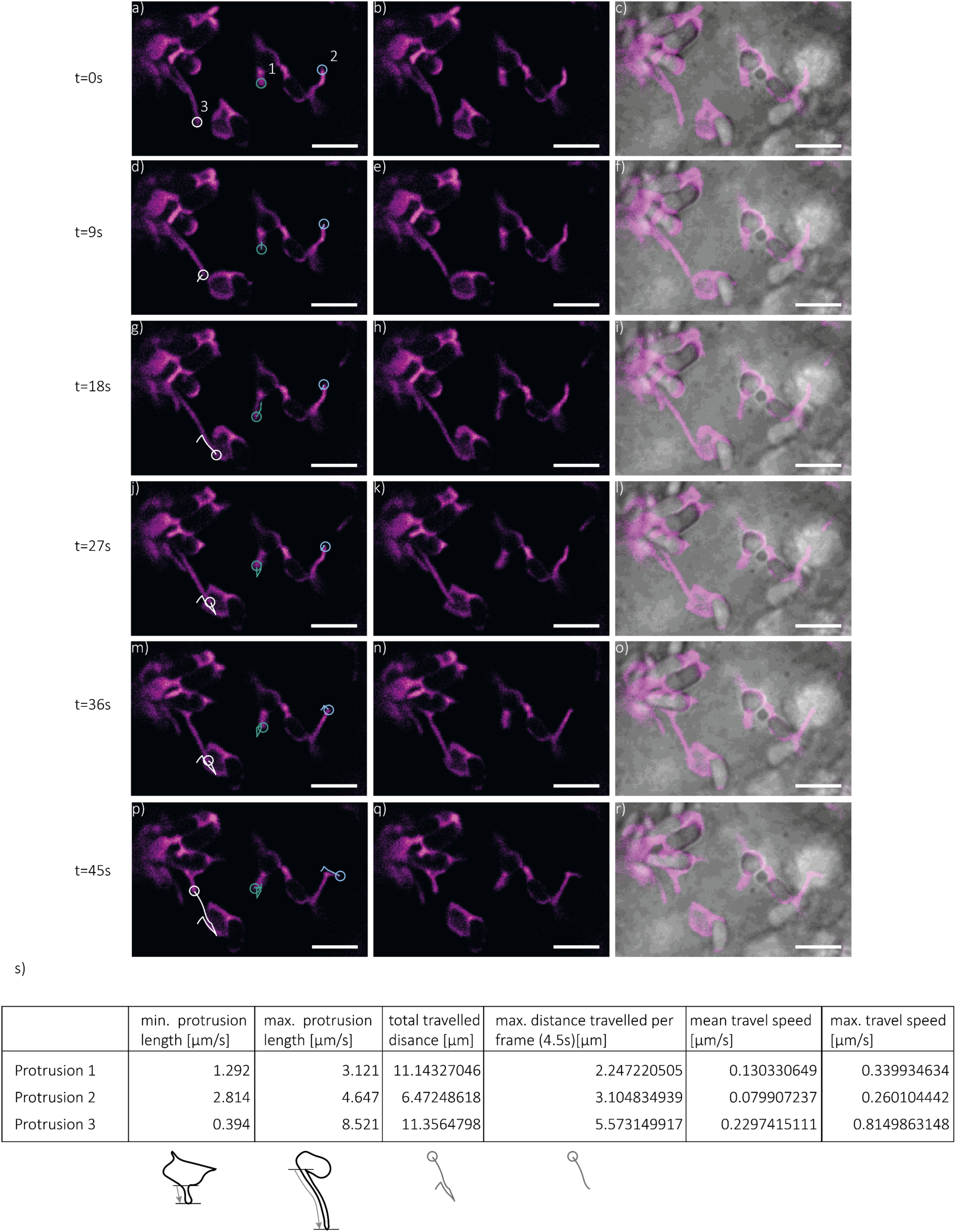
Amyloplast type II protrusions show dynamic movement. (a-r) Confocal laser scanning imaging time series of endosperm sections of developing grain at 16 days after flowering (DAF), in lines stably expressing the amyloplast marker cTPmCherry shown in magenta. Time(t) is indicated in seconds (s). Circles indicate the ends of type II amyloplast protrusions. Lines indicate TrackMate determined paths of amyloplast protrusion ends over the presented frames. Bar = 5 µm. (s) Table listing min. and max. protrusion length, total travelled distance, max. distance travelled per frame, mean travel speed and max. travel speed of protrusions indicated in (a-r) as determined by TrackMate and line measurements as illustrated below, over all frames of the time series presented in (a-r).

### Amyloplast movement in immature *Brachypodium distachyon* endosperm cells is actin dependent

In our time series acquisitions, we observed that the amyloplasts appeared to move in the cell, similar to a bacterial run-and-tumble-type movement (Fig. 7a-f, Video 4). The impression of amyloplast mobility was enhanced by the non-uniform tumbling of individual starch granule containing compartments. Kymograph analysis of amyloplasts in one cell confirmed consistent amyloplast movement in one direction as well as tumbling, changes in movement direction, and the change from tumbling to a more directed type of movement over time (Fig. 7a-g). To test whether the observed amyloplast movement was driven by cytoplasmic streaming, we inhibited cytoplasmic streaming using the actin polymerisation inhibitor, Latrunculin B. Tracking of amyloplasts in cells treated with Latrunculin B showed no major positional changes (Fig. 7h and I, Video 5), while amyloplasts in the control cells moved traceable distances within the same time period.

**Figure 7:**
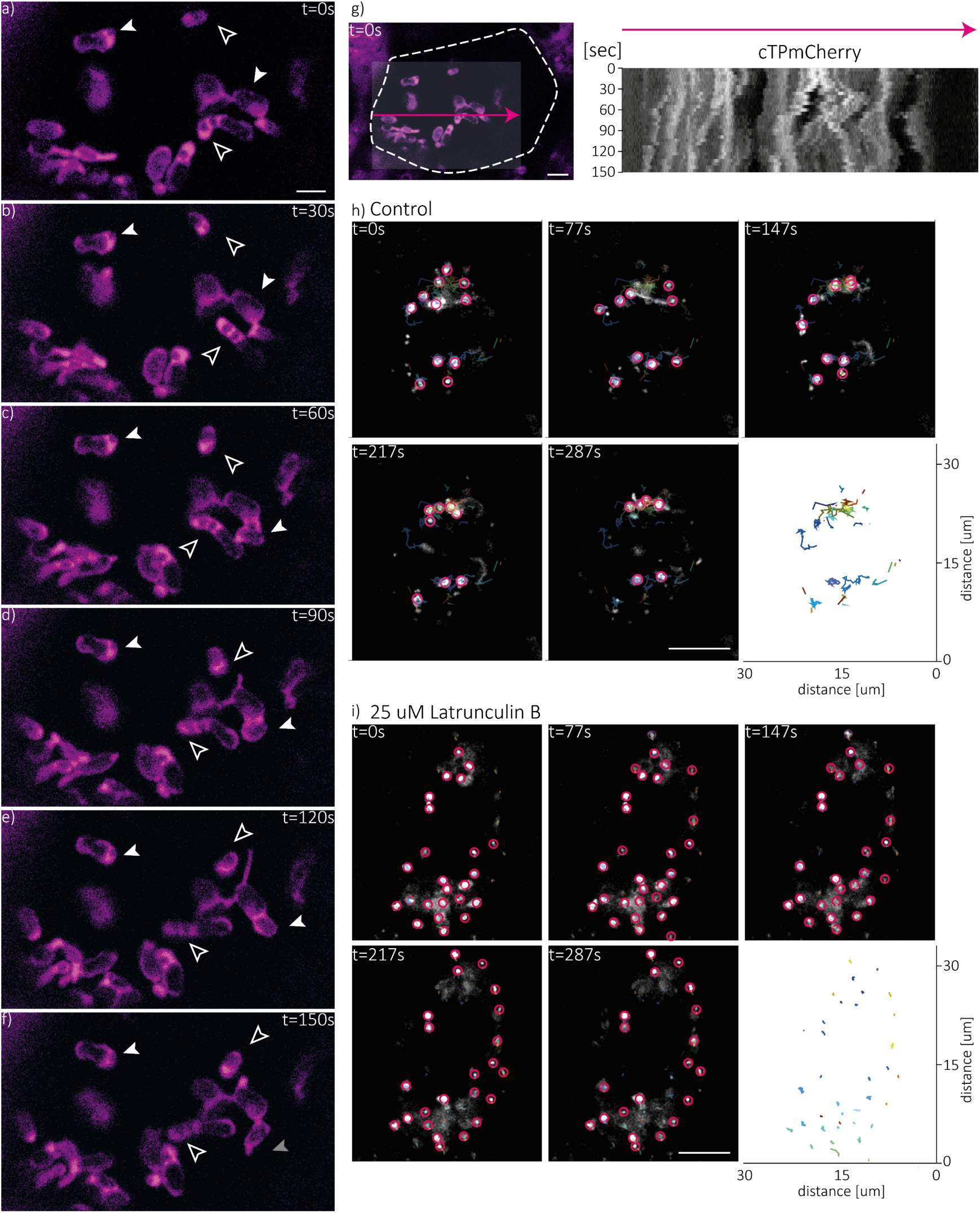
Actin-dependent amyloplast movement in *Brachypodium distachyon* endosperm cells. Amyloplasts were imaged using confocal laser-scanning microscopy in the reporter lines stably overexpressing the amyloplast marker cTPmCherry, shown in magenta. In all panels, time (t) is shown in seconds(s). (a-f) Time series of developing grain at 12 days after flowering (DAF). Hollow arrow indicates directed amyloplast movement, white arrows indicate tumbling, grey arrows indicate shift from tumbling to directed movement. Bar: 5 µm. (g) Kymograph analysis (right) of amyloplasts (magenta) in cell shown in confocal laser scanning image (left). Dashed line indicates cell wall, arrow indicates directionality of kymograph, shaded area represents input into kymograph analysis. Time stamps represented in (a-f) are represented on kymograph y-axis. Bar= 5 µm. (h) Time series of endosperm sections at 16 DAF treated with the control treatment (equivolume of ethanol to Latrunculin B treatment). Time stamps are indicated in seconds (s). Circles indicate amyloplasts tracked with TrackMate, and coloured lines represent tracks. (i) Same as (h), but with sections treated with 25 µM Latrunculin B.

Interestingly, not only whole amyloplasts or amyloplast compartments were mobile, the starch granules within an amyloplast/amyloplast compartment were also able to change their position and orientation relative to one another (Fig. 8, Video 6). In one example, the orientation of the starch granules within one amyloplast compartment showed a maximum displacement of 27° over a timeframe of 350 s (Fig. 8).

**Figure 8:**
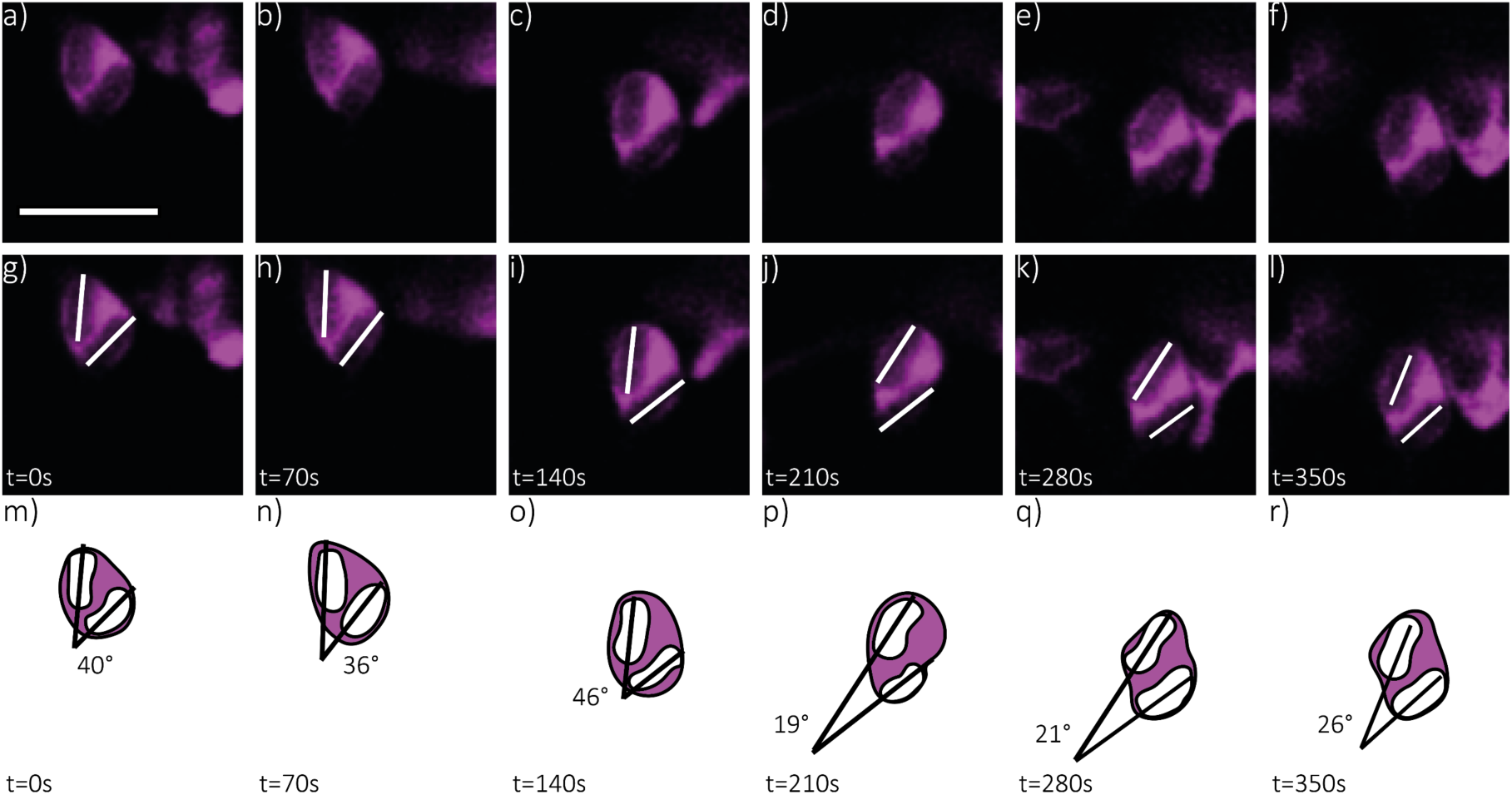
Granule movement within the amyloplast. (a-l) Confocal laser scanning imaging time series of endosperm sections of developing grain at 16 days after flowering (DAF), in lines stably expressing the amyloplast marker cTPmCherry shown in magenta. Time (t) is indicated in seconds (s). Lines indicate starch granule orientation within amyloplast. Bar: 5 µm. (m-r) Schematic representation of starch granule orientation within the amyloplast. Lines represent the angle (in degrees) of starch granules relative to each other.

Overall, our analysis revealed a significant mobility and structural flexibility of endosperm amyloplasts during *Brachypodium* endosperm development. This could potentially allow the starch granules to grow into simple-type starch granules despite there being multiple granules per amyloplast.

## Discussion

### Simple-type granules can form from multiple initiations per amyloplast

Simple starch granules are a common starch granule morphology found not only within grass endosperms (Matsushima et al., 2013), but also more broadly in seed, root and tuber crops, including potato and some yams (Ohad et al., 1971; Matsushima, 2015). Simple granules are typically elliptic, ovoid or round, and were thought to form as a single granule within one amyloplast (Tateoka, 1962; Ohad et al., 1971; Boyer et al., 1977; Echeverria et al., 1988; Kram et al., 1993; Yun and Kawagoe, 2010). This is a key distinction from compound granules, where individual granules have polygonal morphology or flat edges resulting from impaction between multiple granules within the amyloplast (Matsushima et al., 2013; Matsushima et al., 2015).

We made the surprising discovery that endosperm amyloplasts of *Brachypodium* contain multiple granules per amyloplast, despite producing simple rather than compound starch granules (Fig. 1b, 2 and 3). Most starch granules in *Brachypodium* endosperm were initiated during the early stages of grain development, as the number of granules per cell stabilised at 12 DAF (Fig. 1k). However, multiple granules per amyloplast were observed at all stages of grain development examined, including at 9 DAF, suggesting that they are present from early development and retained. Interestingly, we observed a mix of larger and smaller simple-type starch granules within some amyloplasts, while others contained multiple simple-type granules of similar size (Fig. 2, 3 and 4). The heterogeneity in granule size is consistent with a previous study that observed both large and small granules in the *Brachypodium* endosperm (Tanackovic et al., 2014). However, the small granules were present during early grain development, making them distinct to the B-type starch granules in the bimodal granule morphology of the Triticeae, since the small B-type granules in the latter are initiated in separate wave of granule initiation during later grain development (15-20 DAF).

Our findings raise the important question of how the multiple starch granules in *Brachypodium* amyloplasts remain separate and retain simple-type granule morphology, rather than merging into a compound starch granule. Modelling of compound granule formation in rice endosperm suggested that the forces generated from the starch granules and the amyloplast envelope membrane result in the formation of polygonal granulae within a compound granule during endosperm development, resembling a Voronoi diagram (Matsushima et al., 2015). However, flat surfaces from granule impaction were rarely observed in *Brachypodium* starch granules (Fig. 1b). There are reports that the individual granulae of compound granules in rice are separated by membranes derived from the inner amyloplast envelope, as evidenced by the presence of the inner envelope transporter BRITTLE1 in the spaces between the granulae (Yun and Kawagoe, 2010). In the *opaque5* maize mutant, compound granule formation was accompanied by membranes in between the constituent granulae (Myers et al., 2011); in contrast to the compound granules in the wheat *ss4* mutant, where no such separating membranes were observed (Hawkins et al., 2021). We did not observe membranes separating the individual simple granules within *Brachypodium* amyloplasts in our TEM images, suggesting that such internal membranes are unlikely to be the reason why the granules remain separate.

### Amyloplasts in *Brachypodium distachyon* show dynamic changes in structure and stromule formation

Our study demonstrated that *Brachypodium* amyloplasts are pleiomorphic and flexible in shape and size inside endosperm cells throughout grain development (Fig. 2, 3 and 4). Interestingly, we saw that many of the distinct morphological features of the *Brachypodium* endosperm amyloplasts, including the bud-like and vermiform shapes, and protrusions, were still retained when amyloplasts were isolated from their cellular environment (Fig. 3). This suggests that these shapes are maintained by structural features of the amyloplast, like plastid division components or membrane characteristics, rather than external cellular features (e.g., interaction with cytoskeleton).

The structural flexibility and fluidity of the *Brachypodium* endosperm amyloplasts was additionally observed through the formation of two types of amyloplast protrusions. The ‘type I’ protrusions were stable and enabled stromal exchange between amyloplast compartments, as demonstrated through our photobleaching experiments (Fig. 5, Video 1). By contrast, the ‘type II’ protrusions resembled the classical chloroplast stromules, in that they were short-lived, moved and remodelled rapidly and did not connect starch granule containing compartments (Fig. 6, Videos 2 and 3). However, we do not know whether these two types of stromules form via a similar or distinct mechanisms, or whether type I connections form via type II protrusions that have connected to another amyloplast compartment. The formation of both types of stromules are likely to require extensive plastid membrane production, as proposed for stromules in other plastids (Mathur, 2021). Stable connections similar to the type I protrusions were also observed in amyloplast of Arabidopsis ovule integuments, but only in mutants deficient in the plastid division protein MinE (Fujiwara et al., 2024). The *minE* mutant not only formed giant amyloplasts but also extended stable stromules, allowing FtsZ ring assembly and constriction along its length, and the growth of starch granules within the stromule. It is therefore possible that the stable type I protrusions in *Brachypodium* form via an alteration in the plastid division machinery. Indeed, the presence of such stable connections with stromal continuity makes it difficult to define an individual “amyloplast”, as opposed to a series of connected amyloplast compartments. Such interconnected compartments might even be considered as a giant, stretched out plastid. By contrast, type II stromules could form via similar mechanisms to chloroplast stromules, involving extension along microtubules via specific kinesins (Meier et al., 2023).

A variety of different roles have been proposed for plastid stromules, including communication with other organelles, signalling, organelle replication, exchange of metabolites, and defence (Hanson and Hines, 2018; Hanson and Conklin, 2020; Mathur, 2021; Mathur et al., 2023; Fujiwara et al., 2024). In Triticeae endosperm, amyloplast stromules are involved in the formation of bimodal granules, as they facilitate the initiation of B-type starch granules during late grain development (Parker, 1985; Langeveld et al., 2000). This presents a rare example of stromule formation that is triggered at a specific developmental stage and for a defined function. Stromule formation in the Triticeae has also been observed using live-cell imaging of barley and wheat endosperm (Langeveld et al., 2000; Matsushima and Hisano, 2019). We demonstrated that amyloplast stromule formation occurs beyond species that make bimodal granules, suggesting they have other important roles in amyloplast function. It is tempting to speculate that the type I stromules may allow connected compartments to share important metabolites, such as substrates for starch synthesis. It is also possible that they play an important role during amyloplast division. For example, amyloplasts in Arabidopsis ovule integuments were suggested to undergo different modes of replication, including “stromule-mediated replication” where daughter plastids are formed via constricting off a stromule (Fujiwara et al., 2024). However, we note that we did not observe any division event for the *Brachypodium* endosperm amyloplasts during our live-cell imaging.

### Amyloplasts are mobile in the cytoplasmic stream within endosperm cells

Although previous works using live-cell imaging have focused on structure and division dynamics, our work additionally examined movement of amyloplasts within the developing endosperm cells. We observed a bacterial run-and-tumble type movement of the amyloplasts. This movement was inhibited after treatment with Latrunculin B, indicating actin-dependent movement of amyloplasts (Fig. 7). This is similar to chloroplast movement in leaves in response to light (Kasahara et al., 2002; Dwyer and Hangarter, 2022), which is also dependent on actin (Wada and Kong, 2018; Dwyer and Hangarter, 2022; Yamamoto-Negi et al., 2024). Similarly, the non-green plastids of epidermal cells in dark-grown tobacco hypocotyls arrest in mobility with treatment of Latrunculin B (Kwok and Hanson, 2003). Many components involved in actin-driven chloroplast movement have been identified in Arabidopsis, and further investigation is required to determine if they are involved in the amyloplast movement we observed in *Brachypodium*. These include small chloroplast associated actin filaments, F-actin interacting proteins CHUP1 and THRUMIN1, and regulators of actin dynamics in chloroplast movement, KAC1 and KAC2 (Kadota et al., 2009; Oikawa et al.,2003; Oikawa, 2008; Suetsugu et al., 2010; Whippo et al., 2011).

### Consequences of amyloplast structure on simple granule formation

We hypothesise that the observed flexibility and dynamicity of *Brachypodium* amyloplasts could play a central role in facilitating simple, rounded granule formation. When the granules are small, they can likely grow unrestricted in all directions as they are surrounded by stromal space, and the flexibility of the amyloplast membrane can accommodate growing granules. However, like most simple granules, *Brachypodium* starch granules are ovoid (Fig. 1b), suggesting that they undergo anisotropic growth. In potato, a specific isoform of PTST2 (named PTST2b) mediates the anisotropic growth of simple granules, but this isoform is exclusively found in the Solanaceae and not present in *Brachypodium* or cereals (Hochmuth et al., 2025). This implies that there is variation in the mechanism of simple granule formation between species, and there could be other proteins in *Brachypodium* mediating anisotropic granule growth. That said, even in the absence of specific proteins, it is plausible that the amyloplast itself is sufficient to guide directional granule growth. Stretched amyloplast compartments that are interconnected by type I protrusions on either side would have greater membrane tension along the stretched axis, which could restrict growth towards the tensioned faces. Such tension in membranes could increase as the amyloplasts move in the cytoplasmic stream. *Brachypodium* granules also rarely have polygonal morphology or sharp edges, even though multiple granules are present per amyloplast compartment. However, we also observed that individual granules within a moving amyloplast/amyloplast compartment can change their position and orientation relative to each other (Fig. 8). It is plausible that the constant movement, combined with the flexibility of the amyloplast shape, could prevent individual granules from impacting and merging.

Altogether, the dynamic structural changes, stromule formation, and mobility of the *Brachypodium* amyloplasts suggest that these organelles are more structurally complex and pleiomorphic than previously thought. These physical factors likely impact starch granule formation and morphology. Importantly, our study also demonstrates that simple-type starch granules can arise from multiple initiations per amyloplast, challenging the traditional ‘one granule, one amyloplast’ model.

## Materials and Methods

### *Brachypodium distachyon* transformation and copy number analysis

*Agrobacterium*-based transformation of *Brachypodium distachyon* accessions Bd21 was performed according to the Vain et al. (2008) method except with the exclusive use of hygromycin resistance for selection. Briefly, small embryos (<0.3 mm) were isolated from immature seeds and placed on induction media for two weeks to generate compact embryogenic calli (CEC). CEC was sectioned and propagated at two, four, and five weeks of culture at 25°C in the dark, subsequently treated with a seven-minute desiccation, prior to transformation with *Agrobacterium tumefacians* strain AGL1. The T-DNA carried an amyloplast reporter construct (cTPmCherry) as described in Esch et al. (2023), consisting of an N-terminal rice Granule Bound Starch Synthase (GBSS) chloroplast transit peptide (cTP) and a codon optimised mCherry, driven by a maize Ubiquitin promoter (ZmUbi); and the hygromycin phosphotransferase (HPT) selectable marker gene driven by the cauliflower mosaic virus 35S promoter and terminated by the NOS polyA signal. Five weeks after transformation, putatively transformed CEC was transferred to regeneration medium. Three weeks later, plantlets were transferred to containers containing germination medium. Plants were transferred to soil after sufficient growth.

### Growth conditions and sampling

*Brachypodium* wild type (Bd21) and transgenic lines were grown in controlled environment chambers at 60% relative humidity and light intensity of 200 µmol photons m^−2^ s^−1^, with a 16 h light/8 h dark cycle at 24 °C during the day and 18 °C during the night. Tillers were labelled as 0 days after flowering (DAF) as soon as anthers in florets were visible. Developing seeds were collected at 9, 12, 16 and 21 DAF.

### Microscopic analysis of plastid morphology and dynamics

Developing grain of amyloplast reporter lines were harvested at indicated timepoints and hand sectioned into cross sections. Images were acquired immediately after sectioning or amyloplast isolation on a Zeiss LSM880 confocal microscope using a Plan Apo 63.0x/NA 1.4 oil immersion objective (Zeiss). mCherry signal was excited at 561 nm and emission was detected at 562 nm to 623 nm (605 nm).

Fluorescence recovery after photobleaching (FRAP) was carried out on cross sections at 16 DAF seeds using the Zeiss LSM880 confocal microscope with a Plan Apo 63.0x/NA 1.4 oil immersion objective (Zeiss) and a photomultiplier tube (PMT) detector. The laser was set to 561 nm and 2% power for pre- and post-bleach images and 30% for the 5 s bleaching pulse. Pre-bleach (5 images) and post-bleach images (100 images) were captured at either 1.09 s or 0.429 s intervals. The bleaching region was set to the total volume of one starch-containing amyloplast compartment and recovery was measured within this volume as well as connected amyloplast compartments.

Fluorescence correlation spectroscopy (FCS) was used to analyse diffusion speed of mCherry in amyloplasts. FCS was carried out on cross sections of 16 DAF seed using the Stellaris Falcon confocal microscope with an HC Plan Apo 63x/NA 1.2 water immersion objective and a HyDX detector in FCS mode of the LASX software (Leica). Rhodamine B solution (10 nM) was used for calibration. mCherry was excited at 561 nm and detection was set to 600-700 nm. Measurements were taken over 8.3 s and the diffusion coefficient was determined by global fitting of the of the autocorrelation curves with the triple state diffusion model.

The diffusion coefficient of mCherry in amyloplasts was used to determine the mean displacement of an mCherry protein in the amyloplast stroma within the 5 s bleaching pulse used in FRAP.

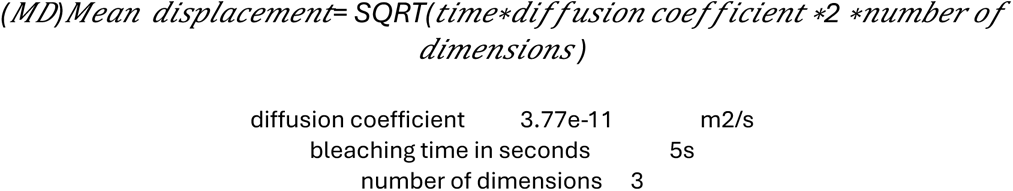

Kymograph of amyloplast movement in cross sections of 12 DAF seed was created based on time series using “Kymograph builder” in Fiji (ImageJ). Tracking of amyloplast movement in cross sections of 16 DAF seeds treated with 25 µM latrunculin B or control solution (containing equivolume of ethanol as the treatment solution) was done in Fiji (ImageJ) using “TrackMate”.

For the analysis of endosperm amyloplast morphology using transmission electron microscopy (TEM): Developing grain (16 DAF) was harvested, the ends were cut off and ca. 2 mm thick cross sections were fixed in 2.5% glutaraldehyde in 0.05 M sodium cacodylate, pH 7.4. Samples were post-fixed in 1% (w/v) osmium tetroxide (OsO4) in 0.05 M sodium cacodylate for 2 h at room temperature, dehydrated in ethanol and infiltrated with LR White resin (Agar Scientific, Stansted, UK), using an EM TP embedding machine (Leica, Milton Keynes, UK). LR White blocks were polymerised at 60°C for 16 h. For transmission electron microscopy (TEM) ultrathin sections (ca. 90 nm) were cut with a diamond knife and placed onto formvar and carbon coated copper grids (EM Resolutions, Sheffield, UK). The sections were stained using 2% (w/v) uranyl acetate for 1 h and 1% (w/v) lead citrate for 1 min, washed in water and air dried. Sections were imaged on a Talos 200C TEM (FEI) at 200 kV and a OneView 4K x 4K camera (Gatan, Warrendale, PA, USA).

Morphology of starch granules in mature grains was examined by cutting a grain in half with razor blade, and imaging the cut surface using a Nova NanoSEM 450 (FEI, Hillsboro, OR, USA).

### Amyloplast isolation

*Brachypodium* endosperm amyloplast isolation was adapted from Denyer and Pike (2008). Endosperm from 16 DAF seeds were harvested into ice-cold 50 mM HEPES-NaOH, pH 7.5 and 0.5 M sorbitol. Plasmolysis took place on ice for 1 h in 50 mM HEPES-NaOH, pH 7.5, 0.8 M sorbitol, 1 mM EDTA, 1 mM KCl, 2 mM MgCl_2_. Plasmolysis solution was replaced once or twice to remove released starch granules. After plasmolysis, the solution was replaced with 50 mM HEPES-NaOH, pH 7.5, 0.8 M sorbitol, 1 mM EDTA, 1 mM KCl, 2 mM MgCl_2_, 1 mM DTT and 1g/L BSA, and the endosperm was gently chopped with a sharp razor blade. Periodically, endosperm pieces were filtered through miracloth and centrifuged at 150*g* at 4°C for 10 min. Amyloplasts in the pellet were resuspended in 50 mM HEPES-NaOH, pH 7.5, 0.8 M sorbitol, 1 mM EDTA, 1 mM KCl, 2 mM MgCl_2_, 1 mM DTT, 1g/L BSA, 50% glycerol and mounted for microscopy.

## Acknowledgements

We thank the John Innes Centre (JIC) Horticultural Services for providing growth facilities and maintenance of plant material, and JIC Bioimaging for providing access to microscopes. We thank Prof Alison Smith (JIC) for helpful advice throughout the project, and Dr. Eva Wegel (JIC) for her expertise on sample preparation and imaging. This work was funded through a Leverhulme Trust Research Project grant RPG-2019-095 (to DS), a John Innes Foundation (JIF) Chris J. Leaver Fellowship (to DS), Gatsby Foundation funding (to MJM), and BBSRC Institute Strategic Programme grants BB/X01097X/1 and BB/X011003/1 (to the John Innes Centre).

## Supplementary Figures and Tables

**Figure S1:**
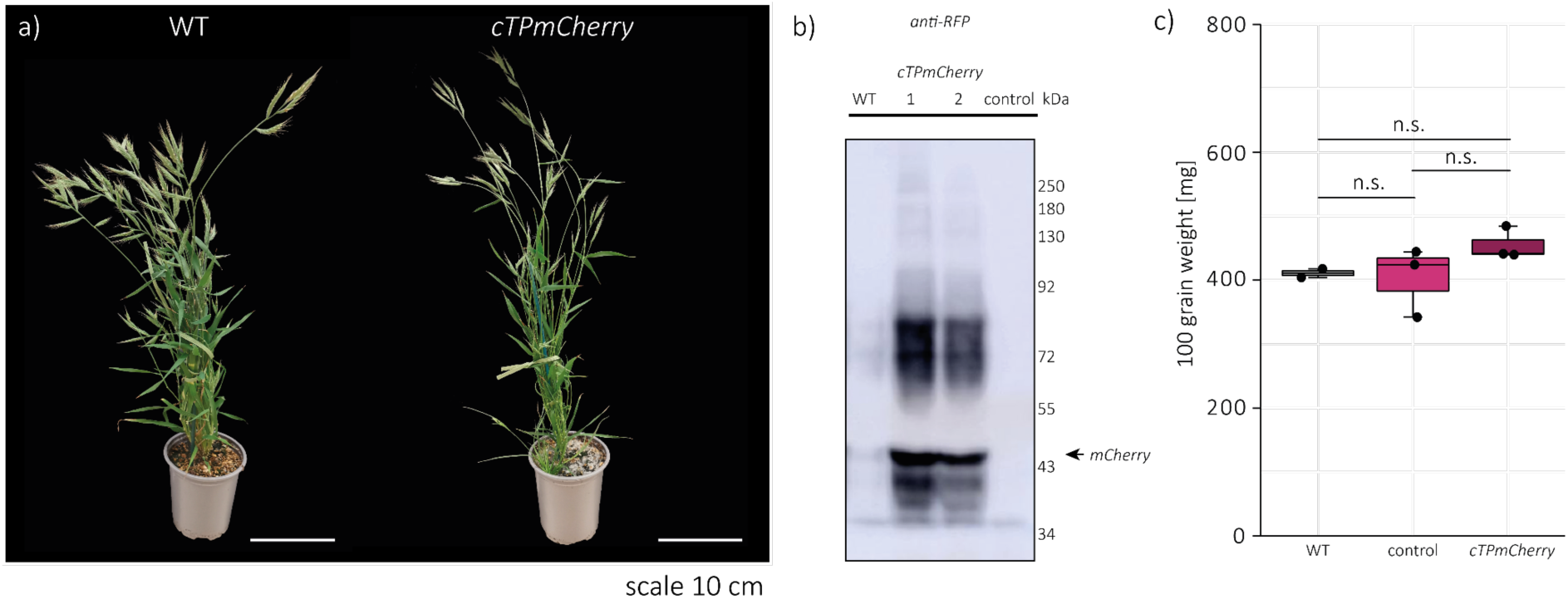
Growth phenotype of amyloplast reporter line. (a) Photograph of 9-week-old Bd21 and Bd21 ZmUbi:cTPmCherry plants. (b) Anti-RFP western blot on total leaf protein of Bd21 (WT), control (plants without the transgene segregating from heterozygous transgenic plants) and Bd21 ZmUbi:cTPmCherry (cTPmCherry) plants (1 C 2). Arrow indicates mCherry protein. Weight in kDa is indicated. (c) Weight of 100 mature grains in mg. 2 biological replicates were included for the WT, and 3 biological replicates for control (plants without the transgene segregating from heterozygous transgenic plants) and Bd21 ZmUbi:cTPmCherry (cTPmCherry) plants. One Way Analysis of Variance did not show any differences between genotypes (p=0.298) with alpha=0.05.

**Figure S2:**
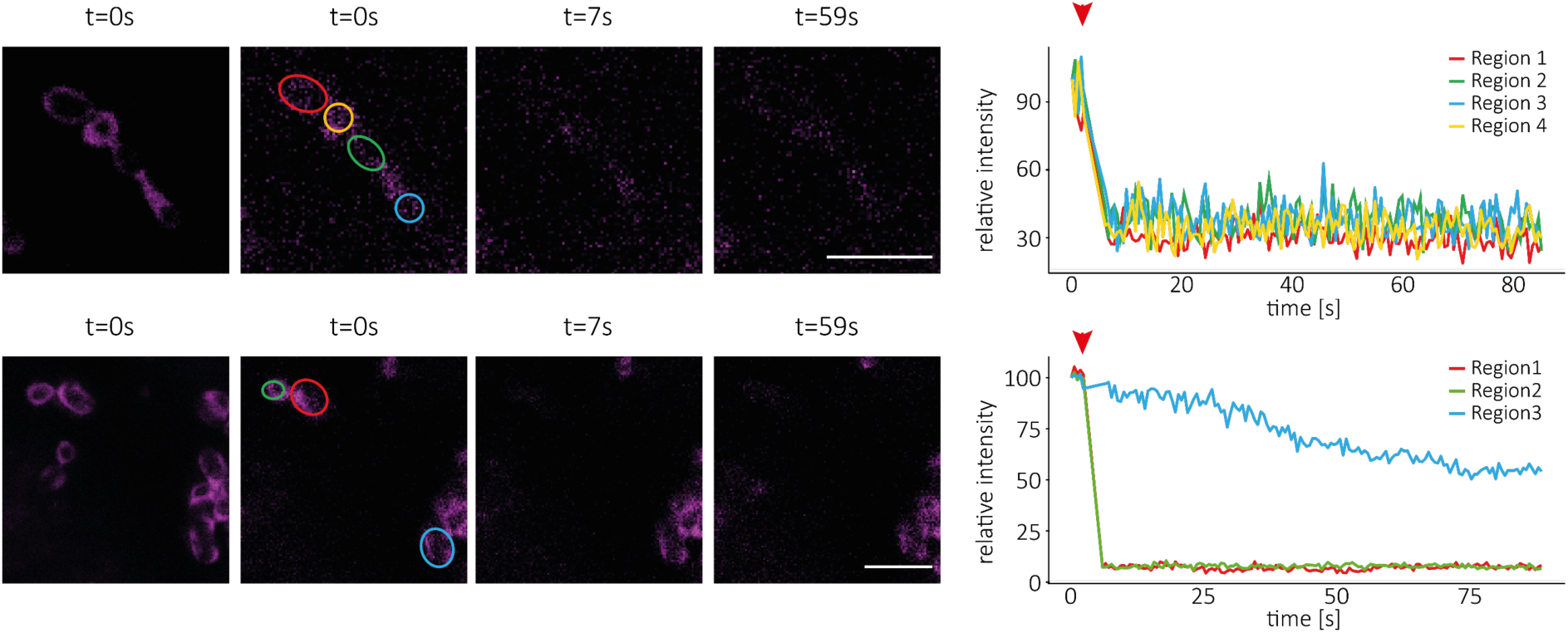
FRAP results for connected amyloplast compartments. (a and c) Confocal laser scanning images of connected amyloplast compartments in endosperm sections of developing grain treated with 25 µM Latrunculin B (a and b) or MS + 3% sucrose (c and d) at 9 days after flowering (DAF), in lines stably overexpressing the amyloplast marker cTPmCherry shown in magenta, before bleaching and after bleaching. Time stamps are indicated in seconds (s). Red circle indicates region of bleaching and fluorescence intensity measurement (Region 1), yellow, green and blue circles indicate regions of fluorescence intensity measurement (Region 2-4). Bar = 5 µm. (b and d) Relative fluorescence intensity after bleaching measured in regions indicated. Red arrow indicates timepoint of bleaching pulse.

## Videos

**Video 1: Amyloplasts in *Brachypodium distachyon* endosperm form protrusions that connect amyloplast compartments.** Confocal laser scanning imaging time series of endosperm sections of developing grain at 16 days after flowering (DAF), in lines stably expressing the amyloplast marker cTPmCherry shown in magenta. Panels show cTPmCherry fluorescence (left) and composite with brightfield (right). Time is shown in seconds (sec). Bar = 10 µm.

**Video 2: Amyloplast type II protrusions show dynamic movement.** Confocal laser scanning imaging time series of endosperm sections of developing grain at 16 days after flowering (DAF), in lines stably expressing the amyloplast marker cTPmCherry shown in magenta. Panels show cTPmCherry fluorescence (left) and composite with brightfield (right). Time is shown in seconds (sec). Bar = 10 µm.

**Video 3: Tracking of amyloplast type II protrusions.** Confocal laser scanning imaging time series of endosperm sections of developing grain at 16 days after flowering (DAF), in lines stably expressing the amyloplast marker cTPmCherry shown in grey. Circles indicate the ends of type II amyloplast protrusions. Lines indicate TrackMate determined paths of amyloplast protrusion ends over the presented frames. Time is shown in seconds (sec). Bar = 5 µm.

**Video 4: Actin-dependent amyloplast movement in *Brachypodium distachyon* endosperm cells.** Confocal laser scanning imaging time series of endosperm sections of developing grain at 12 days after flowering (DAF), in lines stably expressing the amyloplast marker cTPmCherry shown in magenta. Panels show cTPmCherry fluorescence (left) and composite with brightfield (right). Time is shown in seconds (sec). Bar = 10 µm.

**Video 5: Tracking of actin dependent amyloplast movement in the endosperm cell**. Confocal laser scanning imaging time series of endosperm sections of developing grain at 16 days after flowering (DAF), in lines stably expressing the amyloplast marker cTPmCherry shown in grey. Circles indicate the ends of type II amyloplast protrusions. Lines indicate TrackMate determined paths of amyloplast protrusion ends over the presented frames. Time is shown in seconds (sec). Panels show control (ethanol) treatment on the left and Latrunculin B treatment (25 µM) on the right. Bar = 10 µm.

**Video 6: Granule movement within the amyloplast**. Confocal laser scanning imaging time series of endosperm sections of developing grain at 16 days after flowering (DAF), in lines stably expressing the amyloplast marker cTPmCherry shown in magenta. Time is indicated in seconds (sce). Lines indicate starch granule orientation within amyloplast. Bar: 10 µm.

## References

Boyer, C.D., Daniels, R.R., and Shannon, J.C. (1977). Starch granule (amyloplast) development in endosperm of several Zea mays L genotypes affecting kernel polysaccharides. American Journal of Botany 64, 50–56.

Buttrose, M.S. (1960). Submicroscopic development and structure of starch granules in cereal endosperms. Journal of Ultrastructure Research 4, 231–257.

Buttrose, M.S. (1963). Ultrastructure of the developing wheat endosperm. Australian Journal of Biological Sciences 16, 305–317.

Chen, J., Hawkins, E., and Seung, D. (2021). Towards targeted starch modification in plants. Current Opinion in Plant Biology 60, 102013.

Chen, J., Chen, Y., Watson-Lazowski, A., Hawkins, E., Barclay, J.E., Fahy, B., Denley Bowers, R., Corbin, K., Warren, F.J., Blennow, A., Uauy, C., and Seung, D. (2024). Wheat MYOSIN-RESEMBLING CHLOROPLAST PROTEIN controls B-type starch granule initiation timing during endosperm development. Plant Physiol 196, 1980–1996.

Denyer, K., and Pike, M. (2008). Isolation of amyloplasts. Curr Protoc Cell Biol Chapter 3, Unit 3 28.

Dwyer, M.E., and Hangarter, R.P. (2022). Light-induced displacement of PLASTID MOVEMENT IMPAIRED1 precedes light-dependent chloroplast movements. Plant Physiol 189, 1866–1880.

Echeverria, E., Boyer, C.D., Thomas, P.A., Liu, K.C., and Shannon, J.C. (1988). Enzyme activities associated with maize kernel amyloplasts. Plant Physiol 86, 786–792.

Ershov, D., Phan, M.S., Pylvanainen, J.W., Rigaud, S.U., Le Blanc, L., Charles-Orszag, A., Conway, J.R.W., Laine, R.F., Roy, N.H., Bonazzi, D., Dumenil, G., Jacquemet, G., and Tinevez, J.Y. (2022). TrackMate 7: integrating state-of-the-art segmentation algorithms into tracking pipelines. Nat Methods 19, 829–832.

Esch, L., Ngai, Q.Y., Barclay, J.E., and Seung, D. (2022). AtFZL is required for correct starch granule morphology in Arabidopsis chloroplasts. BioRxiv.

Esch, L., Ngai, Q.Y., Barclay, J.E., McNelly, R., Hayta, S., Smedley, M.A., Smith, A.M., and Seung, D. (2023). Increasing amyloplast size in wheat endosperm through mutation of PARC6 affects starch granule morphology. New Phytol 240, 224–241.

Fujiwara, M.T., Yoshioka, Y., Kazama, Y., Hirano, T., Niwa, Y., Moriyama, T., Sato, N., Abe, T., Yoshida, S., and Itoh, R.D. (2024). Principles of amyloplast replication in the ovule integuments of Arabidopsis thaliana. Plant Physiol.

Guillon, F., Larre, C., Petipas, F., Berger, A., Moussawi, J., Rogniaux, H., Santoni, A., Saulnier, L., Jamme, F., Miquel, M., Lepiniec, L., and Dubreucq, B. (2012). A comprehensive overview of grain development in Brachypodium distachyon variety Bd21. J Exp Bot 63, 739–755.

Hanson, M.R., and Hines, K.M. (2018). Stromules: Probing formation and function. Plant Physiology 176, 128–137.

Hanson, M.R., and Conklin, P.L. (2020). Stromules, functional extensions of plastids within the plant cell. Curr Opin Plant Biol 58, 25–32.

Hawkins, E., Chen, J., Watson-Lazowski, A., Ahn-Jarvis, J., Barclay, J.E., Fahy, B., Hartley, M., Warren, F.J., and Seung, D. (2021). STARCH SYNTHASE 4 is required for normal starch granule initiation in amyloplasts of wheat endosperm. New Phytologist 230, 2371–2386.

Hochmuth, A., Carswell, M., Rowland, A., Scarbrough, D., Esch, L., Kamble, N.U., Habig, J.W., and Seung, D. (2025). Distinct effects of PTST2b and MRC on starch granule morphogenesis in potato tubers. Plant Biotechnol J 23, 412–429.

Ilse, T.E., Zhang, H., Heutinck, A., Liu, C., Eicke, S., Sharma, M., Pfister, B., Santelia, D., and Zeeman, S.C. (2025). A point mutation in the photosystem II protein PsbW disrupts thylakoid organization and alters starch granule formation. Plant Physiol 1G8.

Jarvis, P., and López-Juez, E. (2013). Biogenesis and homeostasis of chloroplasts and other plastids. Nature Reviews Molecular Cell Biology 14, 787–802.

Kasahara, M., Kagawa, T., Oikawa, K., Suetsugu, N., Miyao, M., and Wada, M. (2002). Chloroplast avoidance movement reduces photodamage in plants. Nature 420, 829–832.

Kram, A.M., Oostergetel, G.T., and Van Bruggen, E.F.J. (1993). Localization of branching enzyme in potato tuber cells with the use of immunoelectron microscopy. Plant Physiology 101, 237–243.

Kwok, E.Y., and Hanson, M.R. (2003). Microfilaments and microtubules control the morphology and movement of non-green plastids and stromules in Nicotiana tabacum. Plant J 35, 16–26.

Langeveld, S.M.J., Van wijk, R., Stuurman, N., Kijne, J.W., and de Pater, S. (2000). B-type granule containing protrusions and interconnections between amyloplasts in developing wheat endosperm revealed by transmission electron microscopy and GFP expression. Journal of Experimental Botany 51, 1357–1361.

Mathur, J. (2021). Organelle extensions in plant cells. Plant Physiol 185, 593–607.

Mathur, J., Kunjumon, T.K., Mammone, A., and Mathur, N. (2023). Membrane contacts with the endoplasmic reticulum modulate plastid morphology and behaviour. Front Plant Sci 14, 1293906.

Matsushima, R. (2015). Morphological Variations of Starch Grains. In Starch: Metabolism and Structure, Y. Nakamura, ed, pp. 1–451.

Matsushima, R., and Hisano, H. (2019). Imaging Amyloplasts in the Developing Endosperm of Barley and Rice. Scientific Reports 9, 3745.

Matsushima, R., Maekawa, M., and Sakamoto, W. (2015). Geometrical formation of compound starch grains in rice implements Voronoi diagram. Plant and Cell Physiology 56, 2150–2157.

Matsushima, R., Yamashita, J., Kariyama, S., Enomoto, T., and Sakamoto, W. (2013). A phylogenetic re-evaluation of morphological variations of starch grains among Poaceae species. Journal of Applied Glycoscience 60, 131–135.

Meier, N.D., Seward, K., Caplan, J.L., and Dinesh-Kumar, S.P. (2023). Calponin homology domain containing kinesin, KIS1, regulates chloroplast stromule formation and immunity. Sci Adv 9, eadi7407.

Myers, A.M., James, M.G., Lin, Q., Yi, G., Stinard, P.S., Hennen-Bierwagen, T.A., and Becraft, P.W. (2011). Maize opaque5 encodes monogalactosyldiacylglycerol synthase and specifically affects galactolipids necessary for amyloplast and chloroplast function. Plant Cell 23, 2331–2347.

Ohad, I., Friedberg, I., Ne’Eman, Z., and Schramm, M. (1971). Biogenesis and degradation of starch: I. The fate of the amyloplast membranes during maturation and storage of potato tubers. Plant Physiology 47, 465–477.

Parker, M.L. (1985). The relationship between A-type and B-type starch granules in the developing endosperm of wheat. Journal of Cereal Science 3, 271–278.

Seung, D., and Smith, A.M. (2019). Starch granule initiation and morphogenesis – progress in Arabidopsis and cereals. Journal of Experimental Botany 70, 771–784.

Tanackovic, V., Svensson, J.T., Jensen, S.L., Buléon, A., and Blennow, A. (2014). The deposition and characterization of starch in Brachypodium distachyon. Journal of Experimental Botany 65, 5179–5192.

Tateoka, T. (1962). Starch grains of endosperm in grass systematics. Bot. Mag. Tokyo 75, 377–383.

Trafford, K., Haleux, P., Henderson, M., Parker, M., Shirley, N.J., Tucker, M.R., Fincher, G.B., and Burton, R.A. (2013). Grain development in Brachypodium and other grasses: Possible interactions between cell expansion, starch deposition, and cell-wall synthesis. Journal of Experimental Botany 64, 5033–5047.

Vain, P., Worland, B., Thole, V., McKenzie, N., Alves, S.C., Opanowicz, M., Fish, L.J., Bevan, M.W., and Snape, J.W. (2008). Agrobacterium-mediated transformation of the temperate grass Brachypodium distachyon (genotype Bd21) for T-DNA insertional mutagenesis. Plant Biotechnol J 6, 236–245.

Wada, M., and Kong, S.G. (2018). Actin-mediated movement of chloroplasts. J Cell Sci 131.

Wang, H., Huang, Y., Xiao, Q., Huang, X., Li, C., Gao, X., Wang, Q., Xiang, X., Zhu, Y., Wang, J., Wang, W., Larkins, B.A., and Wu, Y. (2020). Carotenoids modulate kernel texture in maize by influencing amyloplast envelope integrity. Nat Commun 11, 5346.

Watson-Lazowski, A., Raven, E., Feike, D., Hill, L., Barclay, J.E., Smith, A.M., and Seung, D. (2022). Loss of PROTEIN TARGETING TO STARCH 2 has variable effects on starch synthesis across organs and species. J Exp Bot 73, 6367–6379.

Yamamoto-Negi, Y., Higa, T., Komatsu, A., Sasaki, K., Ishizaki, K., Nishihama, R., Gotoh, E., Kohchi, T., and Suetsugu, N. (2024). A Kinesin-Like Protein, KAC, is Required for Light-Induced and Actin-Based Chloroplast Movement in Marchantia polymorpha. Plant Cell Physiol 65, 1787–1800.

Yun, M.S., and Kawagoe, Y. (2009). Amyloplast division progresses simultaneously at multiple sites in the endosperm of rice. Plant & cell physiology 50, 1617–1626.

Yun, M.S., and Kawagoe, Y. (2010). Septum formation in amyloplasts produces compound granules in the rice endosperm and is regulated by plastid division proteins. Plant and Cell Physiology 51, 1469–1479.

